# Openness to experience is associated with neural and performance measures of memory in older adults

**DOI:** 10.1101/2022.10.30.514257

**Authors:** Christopher Stolz, Ariane Bulla, Joram Soch, Björn H. Schott, Anni Richter

**Affiliations:** Leibniz Institute for Neurobiology (LIN), Magdeburg, Germany; Institute of Psychology, Otto-von-Guericke University Magdeburg, Magdeburg, Germany; German Center for Neurodegenerative Diseases (DZNE), Göttingen, Germany; Bernstein Center for Computational Neuroscience (BCCN), Berlin, Germany; Department of Psychiatry and Psychotherapy, University Medical Center Göttingen, Göttingen, Germany; Center for Intervention and Research on adaptive and maladaptive brain Circuits underlying mental health (C-I-R-C), Jena-Magdeburg-Halle

**Author notes:** Correspondence Anni Richter, Leibniz Institute for Neurobiology, Department of Behavioral Neurology, Brenneckestraße 6, D-39118 Magdeburg, Germany. Christopher Stolz, Otto-von-Guericke University, Department of Neuropsychology Universitätsplatz 2, D-39106 Magdeburg, Germany.

**Keywords:** openness to experience, episodic memory, aging, fMRI, subsequent memory effect

## Abstract

Age-related decline in episodic memory performance is a well-replicated finding across numerous studies. Recent studies focusing on aging and individual differences found that the Big Five personality trait Openness to Experience is associated with better episodic memory performance in older adults, but the associated neural mechanisms are largely unclear. Here we investigated the relationship between Openness and memory network function in a sample of 352 participants (143 older adults, 50-80 years; 209 young adults, 18-35 years). Participants underwent functional magnetic resonance imaging (fMRI) during a visual memory encoding task. Functional memory brain-network integrity was assessed using the SAME scores (similarity of activations during memory encoding), which reflect the similarity of a participant’s memory network activity compared to prototypical fMRI activity patterns of young adults. Openness was assessed using the NEO Five Factor Inventory (NEO-FFI). Older vs. young adults showed lower memory performance and higher deviation of fMRI activity patterns (i.e., lower SAME scores). Specifically in older adults, high Openness was associated with better memory performance, and mediation analysis showed that this relationship was partially mediated by higher SAME scores. Our results suggest that trait Openness may constitute a protective factor in cognitive aging by better preservation of the brain’s memory network.

## Introduction

Episodic memory is often defined as the ability to encode and recall information of experienced previous events (Baddeley, 2001; Tulving, 2002). A decline in episodic memory performance has been widely characterized as a hallmark symptom of a number of age-related neurological conditions, particularly Alzheimer’s disease (AD; El Haj et al., 2016; Tromp et al., 2015), but it has also been associated with neurocognitive aging in healthy adults (Cansino, 2009; Salthouse, 2010; Grady, 2012). Despite the age-related episodic memory decline, there is considerable inter-individual variability in memory performance in older adults, presumably due to protective factors for age-related neurocognitive declines (Cansino, 2009), such as physical activity (Lee *et al*., 2010; Bherer *et al*., 2013), education (Hendrie *et al*., 2006), and intelligence capacities (Bråthen *et al*., 2021). At personality trait level, the Big Five trait Openness to Experience (hereafter: Openness) has been associated with episodic memory performance (Gregory *et al*., 2010; Terry *et al*., 2013; Curtis *et al*., 2015; Luchetti *et al*., 2016; Sutin *et al*., 2019), with older adults scoring high in this trait showing better memory performance. While the association between Openness and episodic memory performance in older adults is a well-replicated finding, the underlying brain processes are still largely unknown.

In neuroimaging studies, episodic memory encoding and retrieval have been associated with activations of the medial temporal lobe system (i.e., hippocampus, parahippocampal and perirhinal cortex), as well as of prefrontal, parietal, and occipital cortices (Davachi, 2006; Rugg and Vilberg, 2013; Nenert *et al*., 2014). Collectively, these brain regions can be considered to constitute a human memory network (Soch, Richter, Schütze, Kizilirmak, Assmann, Behnisch, *et al*., 2021). Notably, older subjects do not only show reduced activations in this network, particularly in occipital and fusiform cortices, but also reduced deactivations in brain regions of the default mode network (DMN) like the precuneus and posterior cingulate cortex (Maillet and Rajah, 2014; Fjell *et al*., 2016; Damoiseaux, 2017; Soch, Richter, Schütze, Kizilirmak, Assmann, Behnisch, *et al*., 2021; Soch, Richter, Schütze, Kizilirmak, Assmann, Knopf, *et al*., 2021). These memory-related regions are also subject to age-related structural changes in both grey and white matter volumes (Raz and Rodrigue, 2006; Head *et al*., 2008; Fjell and Walhovd, 2010). However, results are mixed regarding the relationship between age-related structural differences and cognitive performance (Salthouse, 2011; Oschwald *et al*., 2019). In a recent study, we showed that chronological age is generally better predicted by structural MRI, whereas functional MRI data, and particularly single-value scores, are superior in predicting individual memory performance in older adults (Soch *et al*., 2022), possibly due to individual differences in compensatory mechanisms to maintain cognitive abilities (Raz and Lindenberger, 2011; Cabeza, Albert, Belleville, *et al*., 2018; Tang *et al*., 2018; Bråthen *et al*., 2021). As such functional age-related differences are often but not always accompanied by cognitive decline (Cansino, 2009; Wong *et al*., 2012; Richter *et al*., 2023), they may also reflect adaptive strategies that are employed as a compensatory mechanism (Grady and Craik, 2000; Cabeza *et al*., 2002; Stern, 2009; Maillet and Rajah, 2014; Cabeza, Albert, Craik, *et al*., 2018). It was proposed that the time point from which on behavioral performance declines during aging strongly relates to an individual’s ability to compensate age-related physiological alterations (e.g., Stern, 2009). While previous behavioral studies indicate that personality facets associated with Openness positively influence the maintenance of higher cognitive functioning including memory (Sharp *et al*., 2010; Ihle *et al*., 2019; Karsazi *et al*., 2021), a possible relationship of Openness with the recruitment of the functional brain networks during successful encoding is yet widely unknown.

To quantify the degree of age-related deviation of fMRI activation patterns from prototypical activity in young adults, single-value scores of activation (and deactivation) patterns during episodic memory encoding have been proposed as a potential tool for the quantification of neurocognitive aging (Düzel *et al*., 2011; Soch, Richter, Schütze, Kizilirmak, Assmann, Behnisch, *et al*., 2021). Düzel et al. (2011) described the FADE score (functional activity deviation during encoding), a measure of deviation from prototypical memory network activations. More recently, Soch, Richter, Schütze, Kizilirmak, Assmann, Behnisch, et al. (2021) introduced the SAME score (similarity of activations during memory encoding), which measures similarity with prototypical brain activity of young adults and reflects both brain activation (i.e. inferior and medial temporal structures, particularly of the parahippocampal cortex, occipital and fusiform cortices) and deactivation patterns (i.e. midline cortical structures). The SAME score correlates positively with memory performance in older adults, that is, higher similarity of encoding-related activations is predictive for better episodic memory performance. This holds for dependent measures (i.e. obtained from the same fMRI task; Soch, Richter, Schütze, Kizilirmak, Assmann, Behnisch, *et al*., 2021) as well as for independent measures (Richter *et al*., 2023) of memory performance. As a reductionist measure, this single-value scores are suitable for investigating correlations with other variables offering a unique possibility to investigate the complex interplay of age-related functional differences, memory decline and personality traits (Allen and DeYoung, 2015; DeYoung and Allen, 2019).

Openness reflects the tendency to be creative, imaginative, curious, and open to new ideas and art (Costa and McCrae, 1989). Highly open individuals have been characterized to show a predisposition towards cognitive flexibility and exploration (DeYoung *et al*., 2005; DeYoung, 2013; DeYoung, 2015). Openness is considered a highly cognitive trait (DeYoung, 2013; DeYoung, 2014) and has been linked to better performance in cognitive abilities such as working memory, inhibitory control, as well as fluid and crystallized intelligence (DeYoung *et al*., 2005; Chamorro-Premuzic and Furnham, 2008). An fMRI study showed that trait Openness was related to working memory accuracy and this effect was fully mediated by activity in left lateral prefrontal and medial frontal cortices (DeYoung *et al*., 2009). In addition to replicating the positive relationship between Openness and intelligence measures, Soubelet & Salthouse (2010) reported that individuals scoring high on trait Openness also showed better episodic memory performance. Large cohort studies examined links between individual differences in Big Five personality traits and age-related episodic memory performance. In a meta-analysis focusing on older adults (>50 years), Luchetti et al. (2016) reported that high Openness, high Conscientiousness, and low Neuroticism were associated with better memory performance and lower memory decline over a four-year longitudinal follow-up. The cross-sectional result pattern was replicated in another sample(Sutin *et al*., 2019), and another longitudinal study showed that individuals scoring high on Openness and low on Neuroticism performed better in an episodic memory task 20 years later(Stephan *et al*., 2020). While there were slightly different correlation patterns of Big Five personality traits and episodic memory performance among those studies, low Neuroticism and high Openness were consistently reported to be associated with better memory performance in older adults. This overall supports the notion of trait Openness as a potential protective factor (and Neuroticism a potential risk factor) for memory function in aging.

While there is evidence supporting a relationship of better episodic memory performance in older adults with both high trait Openness and better-preserved episodic memory brain-network activity, it is yet unclear whether brain activity patterns could serve as a link between individual differences in Openness and memory performance. To this end, the present study aimed to investigate the relationship between Openness, episodic memory performance, and neural episodic memory encoding in young and older adults using personality assessment and fMRI in a visual memory encoding task. Based on previous studies, we hypothesized that trait Openness would be positively related to (1) episodic memory performance and (2) preserved levels of memory brain-network activity (i.e., higher SAME scores) in older adults. Moreover, we also want to answer the question if (3) the relationship between trait Openness and episodic memory performance is mediated by the levels of memory-related brain activity in older adults. While the present study focuses on Openness and memory contrasts, we additionally explored other Big Five personality traits and fMRI novelty contrasts.

## Methods

### Participants

We investigated and combined two previously described study cohorts of 117 (Assmann *et al*., 2021) and 259 (Soch, Richter, Schütze, Kizilirmak, Assmann, Knopf, *et al*., 2021) healthy adults. As we were interested in personality traits, participants were included only if personality questionnaire data were available. These personality attributes have not been analysed or published previously. The total sample of 352 adults was split up into groups of young (n = 209, 18 to 35 years old; age M = 24.30, SD = 3.29) and older adults (n = 143, 50 to 80 years old; age M = 63.80, SD = 6.68). Participants were recruited via flyers at the local academic institutes, advertisement in local newspapers and during public events of the Leibniz Institute for Neurobiology (e.g*., Long Night of the Sciences*). All participants were right-handed, fluent in German language, and did not report any neurological or mental disorder in a standardized neuropsychiatric interview. The local ethics committee of the Faculty of Medicine at the Otto von Guericke University Magdeburg approved the study. All participants signed informed consent in accordance with the Declaration of Helsinki (World Medical Association, 2013) and received financial compensation for participation.

### Visual memory encoding task

In an fMRI session, participants performed an incidental visual memory encoding task (Düzel *et al*., 2018; Assmann *et al*., 2021; Soch, Richter, Schütze, Kizilirmak, Assmann, Knopf, *et al*., 2021). More specifically, participants viewed images of indoor and outdoor scenes and were asked to classify the photos as indoor or outdoor via button press. The task consisted of 132 trials, including unknown indoor and outdoor images (44 trials each; “novel” images) and two images which were pre-familiarized to the participants (one indoor, one outdoor, each presented in 22 trials; “master” images). Each trial started with the presentation of an image for 2500 ms followed by a fixation cross for a variable delay between 500 to 2300 ms in the first cohort (Assmann *et al*., 2021) and 700 to 2650 ms in the second cohort (Soch, Richter, Schütze, Kizilirmak, Assmann, Knopf, *et al*., 2021).

Approximately 70 minutes after the start of the fMRI session, participants underwent a surprise recognition memory test outside the scanner. In this test, the 88 images from the fMRI session (“old”) and 44 at this time unknown images (“new”) were presented. Participants were asked to rate the familiarity of each image on a Likert scale from 1 (“definitely new”) to 5 (“definitely old”). Data were collected using custom code written in Presentation (Version 0.55, Neurobehavioral Systems, www.neurobs.com).

### Memory performance score A’

For the memory test of the pictures shown during fMRI scanning, memory performance was quantified as A-prime (A’), the area under the curve from the receiver-operating characteristic describing the relationship between false alarms (“old” responses to new items) and hits (“old” responses to previously seen items; see Soch, Richter, Schütze, Kizilirmak, Assmann, Behnisch, et al., 2021, Appendix B).

More precisely, let *o*_1_, …, *o*_5_ and *n*_1_, …, *n*_5_ be the numbers of old stimuli and new stimuli, respectively, rated during retrieval as 1 (“definitely new”) to 5 (“definitely old”). Then, hit rates and false alarm (FA) rates as functions of a threshold *t* ∈ {0,1, …, 5} are given as the proportions of old stimuli and new stimuli, respectively, rated higher than *t*:

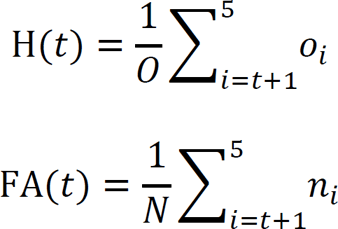

where *o*_1_ + ⋯ + *o*_5_ and *N* = *n*_1_ + ⋯ + *n*_5_. Note that H(0) = FA(0) = 1 and H(5) = FA(5) = 0. Consider the hit rate as a function of the FA rate:

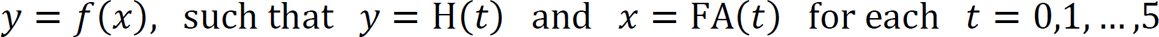

Then, the area under the ROC curve is given as the integral of this function from 0 to 1:

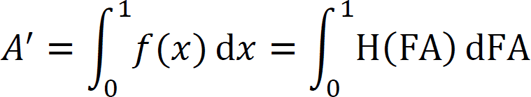

This quantity is referred to as “A-prime” and serves as a measure for memory performance (Soch, Richter, Schütze, Kizilirmak, Assmann, Behnisch, et al., 2021). A’ ranges from 0 to 1 with A’ = 0.5 indicating random guessing and A’ = 1 perfect performance

### fMRI data acquisition and preprocessing

The fMRI data were assessed using two Siemens 3T MR tomographs (Verio and Skyra). Functional MRI was assessed as 206 T2*-weighted echo-planar images (EPIs; first cohort: TR = 2.4 s, TE = 30 ms, flip-α = 80°; 40 slices, 104 × 104 in-plane resolution, voxel size = 2 × 2 × 3 mm, for details see Assmann et al., 2021; second cohort: TR = 2.58 s, TE = 30 ms, flip-α = 80°; 47 slices, 64 × 64 in-plane resolution, voxel size = 3.5 × 3.5 × 3.5 mm, for details see Soch, Richter, Schütze, Kizilirmak, Assmann, Knopf, et al., 2021) in interleaved-ascending slice order (1, 3, … 47, 2, 4, …, 46). Structural MRI was acquired as a T1-weighted MPRAGE image (TR = 2.5 s, TE = 4.37 ms, flip-α= 7°; 192 slices, 256 ×256 in-plane resolution, voxel size = 1 × 1 × 1 mm) for later co-registration.

Data preprocessing and analysis were performed using Statistical Parametric Mapping (SPM12; Wellcome Trust Center for Neuroimaging, University College London, London, UK; https://www.fil.ion.ucl.ac.uk/spm/software/spm12/). EPIs were corrected for acquisition delay, head motion, and in the second cohort additionally for magnetic field inhomogeneities using voxel-displacement maps derived from the field maps. Finally, unwarped EPIs were normalized to a common voxel size of 3 × 3 × 3 mm and eventually spatially smoothed using an isotropic Gaussian kernel of 6 mm at FWHM.

### Extraction of single-value FADE and SAME fMRI scores

FADE and SAME scores were calculated following the protocol of previous studies (Soch, Richter, Schütze, Kizilirmak, Assmann, Behnisch, *et al*., 2021; Richter *et al*., 2023). First, we generated single-subject contrast images representing effects of novelty processing (contrasting novel with master images) and subsequent memory effects (parametrically modulating the BOLD response to novel images as a function of arcsine-transformed subject’s responses ranging from 1 to 5 in the subsequent recognition memory test). Second, we computed a reference map showing significant activations (and, for the SAME score, additionally significant deactivations) on each of the two fMRI contrasts (i.e. novelty processing or subsequent memory) within young adults. Third, we calculated summary statistics for every participant and fMRI contrast: the FADE score (indicating the amount of deviation of activations from young subjects) and the SAME score (indicating the amount of similarities of activations and deactivations with young subjects). The SAME score additionally accounts for the inter-individual variability within the reference sample of young subjects via dividing by their standard deviation.

More precisely, let *J*_+_ be the set of voxels showing a positive effect in young subjects at an *a priori* defined significance level (here: *p* < 0.05, FWE-corrected, extent threshold k = 10 voxels), and let *t_ij_* be the t-value of the *i*--th older subject in the *j*-th voxel on the same contrast. Then, the FADE score of this subject is given by

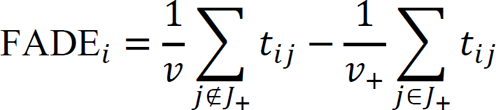

where *v*_+_ and *v* is the number of voxels inside and outside *J*_+_, respectively (Soch, Richter, Schütze, Kizilirmak, Assmann, Behnisch, *et al*., 2021). A larger FADE score signifies higher deviation of an older adult’s memory – or novelty – response from the prototypical response seen in young adults.

Now consider *J*_−_, the set of voxels showing a negative effect in young subjects at a given significance level. Furthermore, let 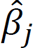 be the average contrast estimate in young subjects, let 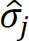 be the standard deviation of young subjects on a contrast at the *j*-th voxel, and let 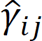 be the contrast estimate of the *i*--th older subject at the *j*-th voxel. Then, the SAME score is given by

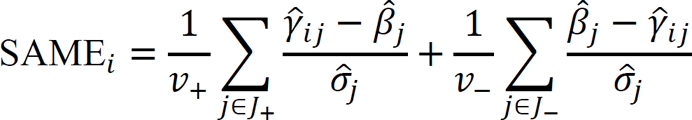

where *v*_+_ and *v*_−_ are the numbers of voxels in *J*_+_ and *J*_−_, respectively (Soch, Richter, Schütze, Kizilirmak, Assmann, Behnisch, *et al*., 2021).

Hereafter, we refer to the scores as follows:

- FADE score computed from the novelty contrast: FADE novelty score;
- SAME score computed from the novelty contrast: SAME novelty score;
- FADE score computed from the memory contrast: FADE memory score;
- SAME score computed from the memory contrast: SAME memory score.

### Assessment of trait Openness to Experience

Openness which reflects self-reports of being imaginative, curious, and having a wide range of interests was assessed using the German version (Borkenau and Ostendorf, 1993; Körner *et al*., 2008) of the NEO-Five Factor Inventory (NEO-FFI; Costa and McCrae, 1989), a shortened version of the NEO-PI-R (Costa and McCrae, 1992). The NEO-FFI is a 60-item questionnaire (12 items for each scale) that measures the Big Five personality traits Neuroticism, Extraversion, Openness, Agreeableness, and Conscientiousness. The items are rated on a five-point Likert scale (strongly disagree to strongly agree). The internal consistency of the scales ranged between a Cronbach’s alpha of 0.71 and 0.85 (Borkenau and Ostendorf, 1993; Schmitz *et al*., 2001; Körner *et al*., 2008).

### Statistical analysis

Data were analyzed using custom code written in MATLAB R2022a and IBM® SPSS® Statistics, Version 28. Comparisons of mean A’, FADE/SAME novelty and memory scores and personality scores between young vs. older adults were performed using *t* tests. For nominally-scale variables (gender and education), the χ^2^-test for independence was used. All reported correlation coefficients were Pearson’s *r* if not stated otherwise.

To examine the independent effects of Openness on A’ and the FADE/SAME memory scores in older adults, multiple regression analyses were calculated. Since previous studies reported substantial covariations of Openness with demographic variables and a range of cognitive functions (DeYoung *et al*., 2005; Gregory *et al*., 2010; Soubelet and Salthouse, 2010), we included age, gender (male vs. female), education (with vs. without German university entrance diploma, “Abitur”), and crystallized intelligence measured with the Multiple-Choice Vocabulary Intelligence Test (Mehrfachwahl-Wortschatz-Intelligenztest B [MWT-B]; Lehrl, 1999) as explanatory variables in the analyses.

Mediation models with Openness as independent variable, A’ as dependent variable, and FADE/SAME memory scores as mediator variable were calculated using the *lavaan* package (Rosseel, 2012) in R (R Core Team, 2013). The reported path estimates for the mediation model were standardized estimates using all path information (Std.all). The reported effect sizes were standardized beta coefficients (β) and Cohen’s *d*. To control for covariations with demographics and crystallized intelligence, the analyses described above were also conducted with adjusted variables. Therefore, multiple regression analyses were run including age, gender, education, and MWT-B scores on Openness, A’, FADE memory score, and SAME memory score and residuals were saved for further mediation analyses.

As an exploratory analysis, we computed voxel-wise regressions of the fMRI subsequent memory contrast with Openness in older adults. Results are reported at *p*cluster < 0.050 using family-wise error rate (FWE) cluster-level correction and an uncorrected cluster-forming threshold of *p*voxel < 0.001 (Eklund *et al*., 2016).

## Results

### Demographic information and trait Openness

Young and older adults did not differ significantly with respect to gender ratio (Table 1). As reported previously, older vs. young adults less frequently had an university entrance diploma (education, “Abitur”), most likely due to historical differences in educational systems (for a detailed discussion, see Soch, Richter, Schütze, Kizilirmak, Assmann, Behnisch, *et al*., 2021). Using the MWT-B (Lehrl, 1999), a screening of crystallized intelligence, we found higher MWT-B scores in older vs. young participants^11^. In the NEO-FFI, older vs. young adults showed lower scores for Openness (*t*(350) = -3.97, *p* < 0.001, *d* = -0.43; Figure 1), Neuroticism (*t*(350) = -2.28, *p* = 0.024, *d* = -0.25), and Extraversion (*t*(350) = -4.16, *p* < 0.001, *d* = -0.45), and higher scores for Conscientiousness (*t*(350) = 2.94, *p* < 0.001, *d* = 0.25). There was no significant group difference for Agreeableness (*t*(350) = -1.08, *p* = 0.283, *d* = -0.25, Table 1).

**Table 1.**
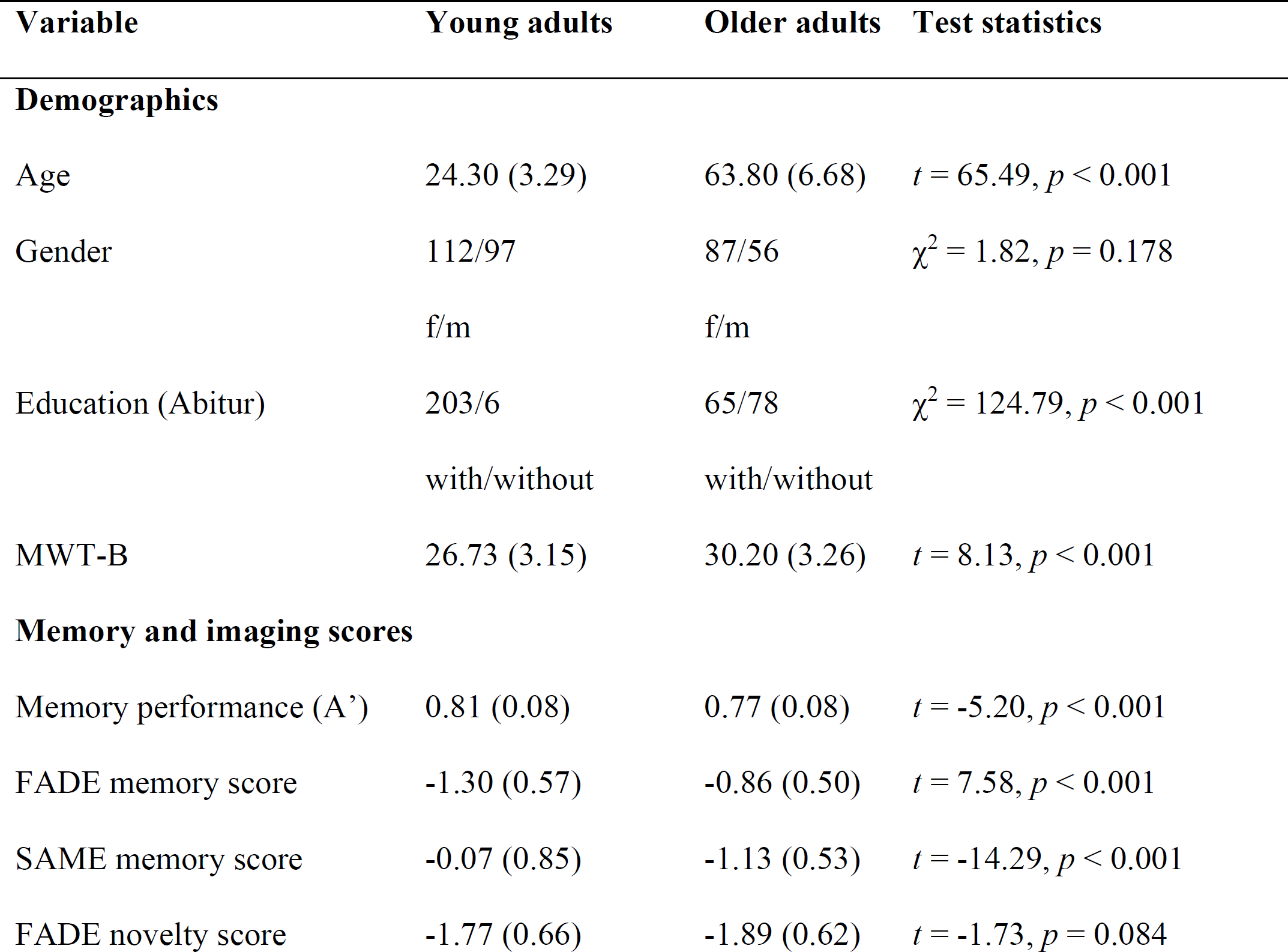
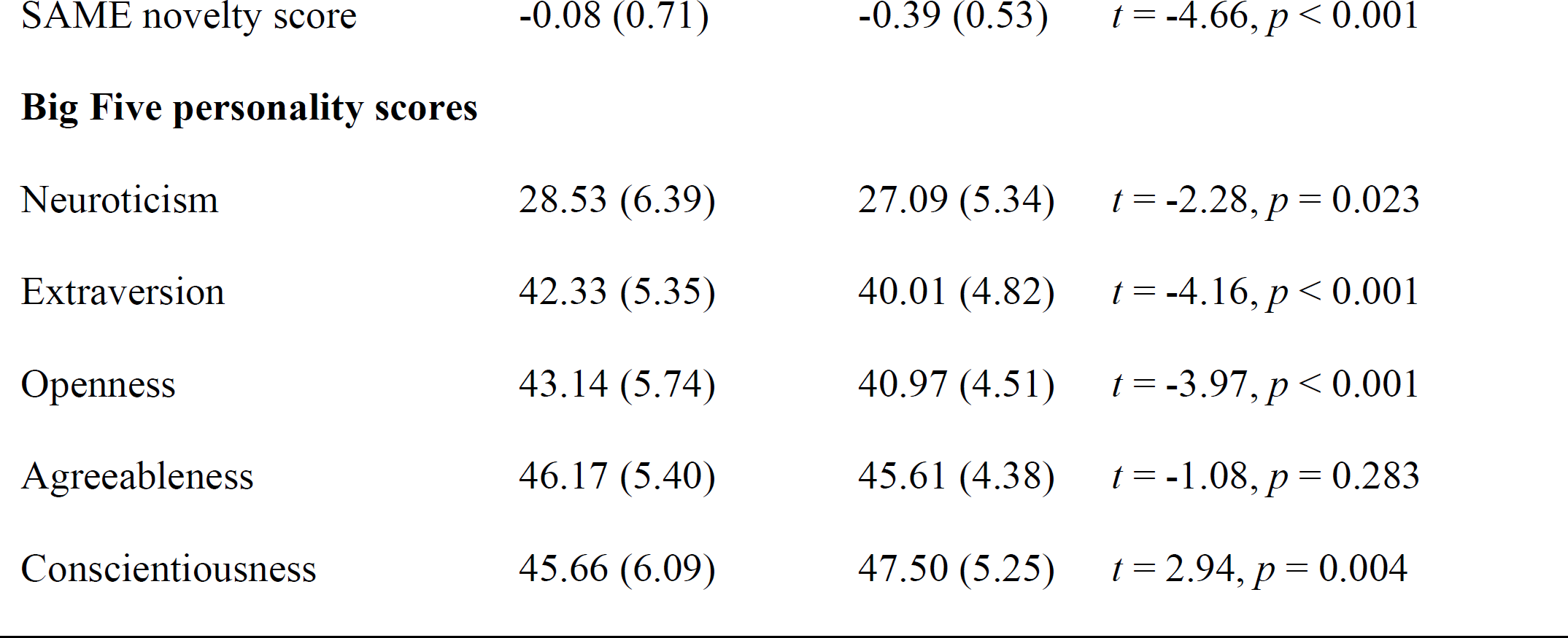
Demographic information and mean (SD) memory, imaging and personality trait scores in the young vs. older adult groups <colcnt=1>

| Variable | Young adults | Older adults | Test statistics |
| --- | --- | --- | --- |
| **Demographics** |  |  |  |
| Age | 24.30 (3.29) | 63.80 (6.68) | $t = 65.49$, $p < 0.001$ |
| Gender | 112/97<br>f/m | 87/56<br>f/m | $\chi^2 = 1.82$, $p = 0.178$ |
| Education (Abitur) | 203/6<br>with/without | 65/78<br>with/without | $\chi^2 = 124.79$, $p < 0.001$ |
| MWT-B | 26.73 (3.15) | 30.20 (3.26) | $t = 8.13$, $p < 0.001$ |
| **Memory and imaging scores** |  |  |  |
| Memory performance (A') | 0.81 (0.08) | 0.77 (0.08) | $t = -5.20$, $p < 0.001$ |
| FADE memory score | -1.30 (0.57) | -0.86 (0.50) | $t = 7.58$, $p < 0.001$ |
| SAME memory score | -0.07 (0.85) | -1.13 (0.53) | $t = -14.29$, $p < 0.001$ |
| FADE novelty score | -1.77 (0.66) | -1.89 (0.62) | $t = -1.73$, $p = 0.084$ |
| SAME novelty score | -0.08 (0.71) | -0.39 (0.53) | $t = -4.66, p < 0.001$ |
| Big Five personality scores |  |  |  |
| Neuroticism | 28.53 (6.39) | 27.09 (5.34) | $t = -2.28, p = 0.023$ |
| Extraversion | 42.33 (5.35) | 40.01 (4.82) | $t = -4.16, p < 0.001$ |
| Openness | 43.14 (5.74) | 40.97 (4.51) | $t = -3.97, p < 0.001$ |
| Agreeableness | 46.17 (5.40) | 45.61 (4.38) | $t = -1.08, p = 0.283$ |
| Conscientiousness | 45.66 (6.09) | 47.50 (5.25) | $t = 2.94, p = 0.004$ |

**Figure 1.**
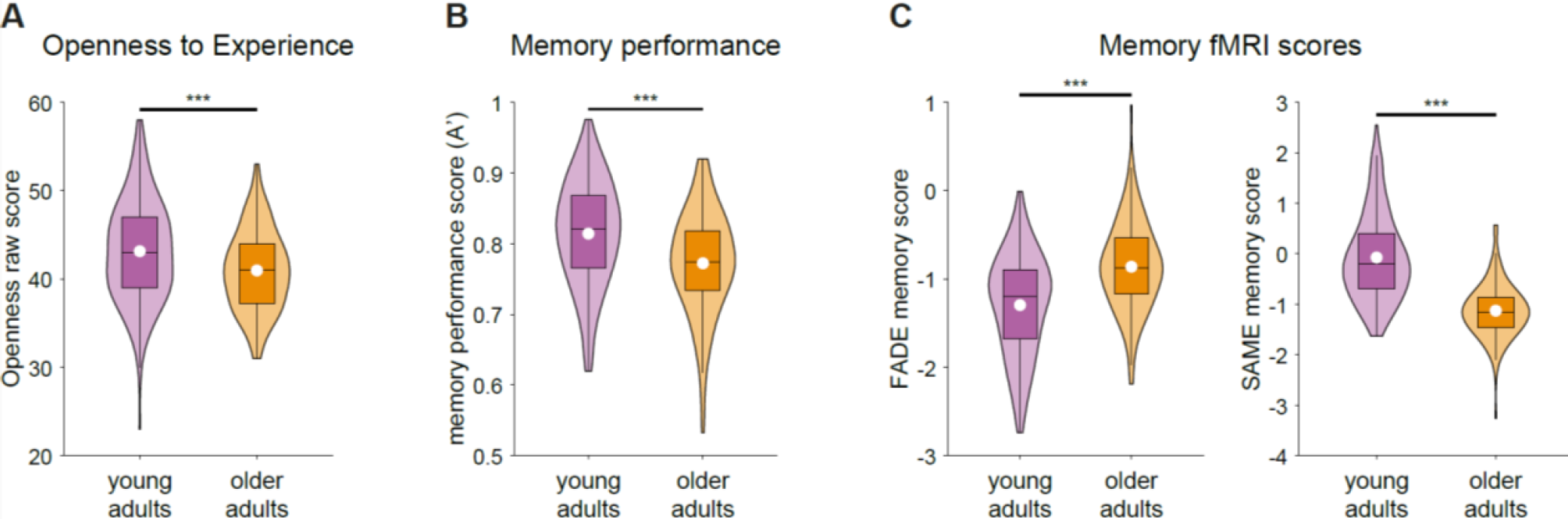
Violin plots of the (A) Openness raw scores, (B) memory performance, and (C) FADE/SAME memory scores in young vs. older adults. The white dot indicates the mean value and the box outlines the interquartile range and median. \*\*\**p* < 0.001<colcnt=1>

### Age differences for memory performance and FADE/SAME scores

A’ was significantly lower in older (M = 0.77, SD = 0.73) vs. young adults (M = 0.81, SD = 0.75; *t*(350) = -5.20, *p* < 0.001, *d* = -0.57; Table 1 and Figure 1). As shown in Figure 1, there was a significant difference between older vs. young adults in the FADE (young: M = -1.30, SD = 0.57; older: M = -0.86, SD = 0.50; *t*(350) = 7.38, *p* < 0.001, *d* = 0.80) and SAME memory scores (young: -0.07, SD = 0.85; older: M = -1.13, SD = 0.53; *t*(350) = -13.17, *p* < 0.001, *d* = - 1.43). Taken together, this suggests a higher deviation (FADE) and lower similarity (SAME) of the memory brain-network activity in older adults compared to the reference of prototypical young adult activity with a particularly large effect size for the SAME score. While the SAME novelty score was significantly different between the age groups (young: -0.08, SD = 0.71; older: M = -0.39, SD = 0.54; *t*(350) = -4.43, *p* < 0.001, *d* = -0.48), there was no significant group difference in the FADE novelty score (young: M = -1.77, SD = 0.66; older: M = -1.89, SD = 0.62; *t*(350) = -1.73, *p* = 0.084, *d* = -0.19). This might be due to differences in the underlying brain networks and their association with aging (e.g., a rather specific association of the FADE novelty score with MTL regions, vs. more widespread associations for all other scores, for illustration see Richter et al., 2023, Figure S2, and for a detailed discussion, see Soch, Richter, Schütze, Kizilirmak, Assmann, Behnisch, et al., 2021).

#### Correlations of memory performance and imaging scores with big five personality traits

While we observed significant correlations between the FADE/SAME memory scores with A’ in both age groups, personality scores particularly correlated with memory performance and fMRI scores in older adults (Table 2). Largest correlations were found for Openness with A’ (*r* = 0.27, *p* = 0.001), FADE (*r* = -0.25, *p* = 0.002) and SAME memory scores (*r* = 0.29, *p* < 0.001), suggesting that older adults scoring high on Openness show better memory performance and more prototypical memory network activity (Figure 2). In the exploratory analysis for the other personality traits, a similar pattern was observed for Extraversion (Table 2). For exploratory correlations between the single NEO-FFI items of Openness, A’, and the FADE/SAME memory scores in older adults, see supplementary Material A. The largest correlation between Openness and A’ was found for “Philosophical discussions”, which belongs to the NEO-PI-R Openness facet “Ideas”. For FADE/SAME memory scores the largest correlations with Openness were found for “Wave of excitement to art/literature”, which belongs to the NEO-PI-R Openness facet “Aesthetics” (Costa and McCrae, 1992; Ostendorf and Angleitner, 2004).

**Table.** Correlation coefficients between the Big Five personality traits and the memory performance (A’) and fMRI scores for the young and older adults

|  |  | Neur. | Extr. | Open. | Agre. | Cons. | A' |
| --- | --- | --- | --- | --- | --- | --- | --- |
| Young | A' | -0.07 | 0.17** | 0.09 | 0.14* | 0.04 | 1 |
| adults | FADE memory score | -0.04 | -0.07 | -0.03 | -0.14 | -0.04 | -0.27** |
|  | SAME memory score | -0.03 | 0.05 | 0.04 | 0.10 | -0.01 | 0.39** |
|  | FADE novelty score | -0.09 | 0.11 | 0.07 | 0.06 | 0.03 | 0.09 |
|  | SAME novelty score | 0.04 | -0.06 | -0.09 | -0.12 | -0.08 | 0.02 |
| Older | A' | -0.14 | 0.25** | 0.27** | 0.01 | -0.02 | 1 |
| adults | FADE memory score | 0.01 | -0.17* | -0.25** | 0.11 | 0.10 | -0.33** |
|  | SAME memory score | 0.01 | 0.17* | 0.29** | -0.07 | -0.13 | 0.41** |
|  | FADE novelty score | -0.04 | 0.04 | 0.12 | 0.15 | -0.07 | -0.04 |
|  | SAME novelty score | -0.01 | 0.13 | -0.09 | -0.03 | 0.15 | 0.24** |

*Notes.* Neur. = Neuroticism, Extr. = Extraversion, Open. = Openness to Experience, AGRE. = Agreeableness, Cons. = Conscientiousness. \*\* $p < 0.01$ , \* $p < 0.05$

**Figure 2.**
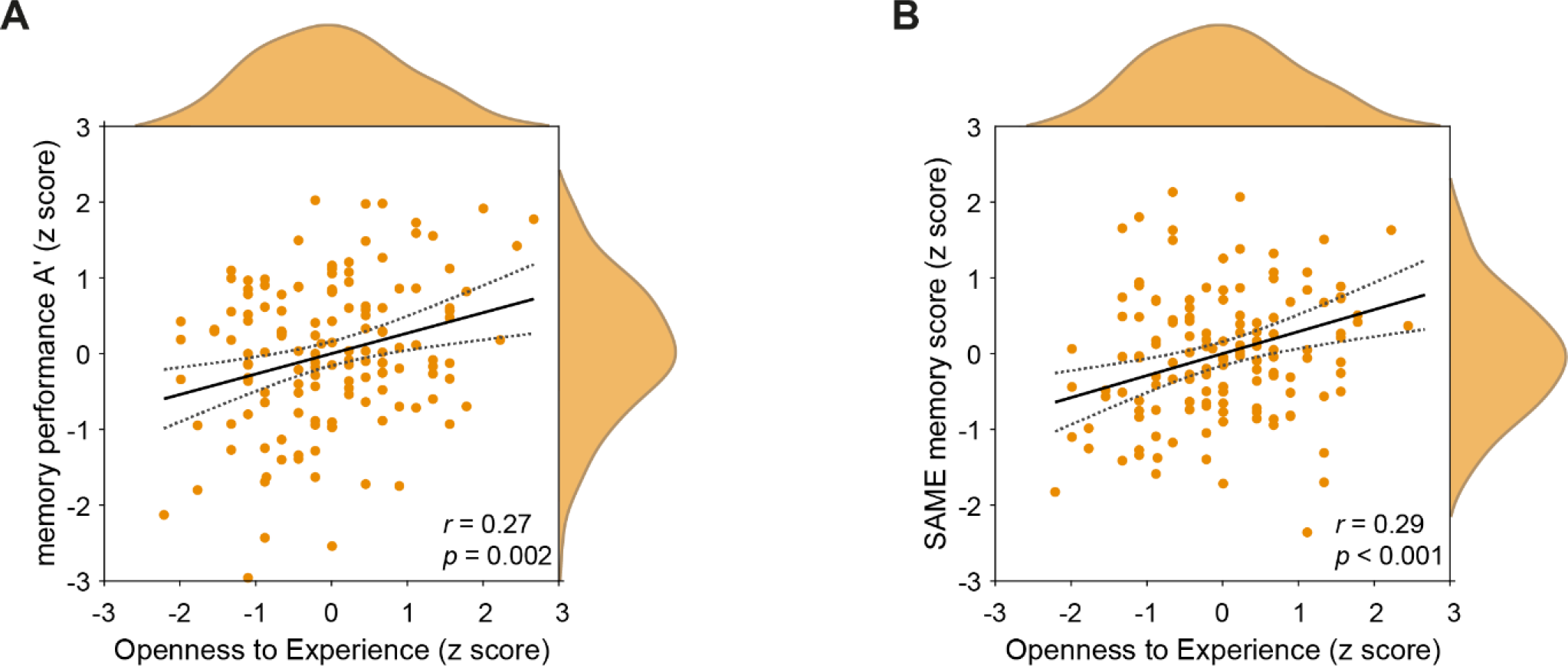
Scatter plots including 95% confidence intervals and the data distribution for the correlations of Openness (here shown as z scores) with (A) the memory performance and (B) the SAME memory score within the group of older adults.

### Trait Openness within the group of older adults

As Openness was correlated with A’ and the FADE/SAME memory scores in older adults only, further analyses are focused on this group. In older adults, Openness was significantly positively correlated with education (point-biserial *r* = 0.22, *p* = 0.008) and MWT-B test results (*r* = 0.18, *p* = 0.036), but not with age (*r* = 0.04, *p* = 0.658) or gender (point-biserial *r* = -0.04, *p* = 0.656). Multiple regression analyses revealed that there was a substantial direct effect of Openness as predictor on A’, FADE memory score and SAME memory score as criterion, independently of the demographic variables (age, gender, and education) and MWT-B scores (Table 3).

**Table 3.** Multiple regression analysis of the direct effects of Openness on memory performance and FADE/SAME memory scores controlled for demographic variables (age, gender, and education) and crystallized intelligence (MWT-B scores) in older adults.

|  | A' |  | FADE memory score |  | SAME memory score |  |
| --- | --- | --- | --- | --- | --- | --- |
| | $\beta$ | $p$ | $\beta$ | $p$ | $\beta$ | $p$ |
| Openness | 0.24** | 0.005 | -0.22* | 0.009 | 0.26** | 0.002 |
| Age | -0.10 | 0.250 | 0.17* | 0.048 | -0.09 | 0.297 |
| Gender | 0.03 | 0.730 | -0.07 | 0.415 | 0.15 | 0.075 |
| Education | 0.09 | 0.321 | -0.11 | 0.213 | 0.18* | 0.048 |
| MWT-B | 0.10 | 0.288 | -0.09 | 0.351 | 0.03 | 0.707 |

*Notes.* One participant was excluded from all analyses involving the MWT-B due to missing value. \*\* $p < 0.01$ , \* $p < 0.05$

As shown in Table 4, the mediation analysis with the FADE memory score as mediator on the relationship between Openness as independent variable and A’ as dependent variable revealed a partial mediation effect (total effect: β = 0.27, *p* = 0.001; indirect effect: β = 0.07, *p* = 0.020). This suggests that the FADE memory score mediated 26.29% of the total effect between Openness and A’.

**Table 4.** Mediation analysis within the group of older adults with FADE/SAME memory scores as mediators of the association between Openness and episodic memory performance (A’).

| FADE memory score | $\beta$ | $P$ |
| --- | --- | --- |
| Total effect: Openness → A' | 0.27 | 0.001 |
| Direct effect: FADE memory score → A' | -0.25 | 0.002 |
| Direct effect: Openness → A' | 0.20 | 0.013 |
| Indirect effect: Openness → FADE memory score → A' | 0.07 (26.29%) | 0.020 |
| **SAME memory score** | $\beta$ | *P* |
| Total effect: Openness → A' | 0.27 | 0.001 |
| Direct effect: SAME memory score → A' | 0.37 | < 0.001 |
| Direct effect: Openness → A' | 0.16 | 0.036 |
| Indirect effect: Openness → SAME memory score → A' | 0.11 (40.74%) | 0.004 |

The mediation analysis with the SAME memory score as mediator on the relationship between Openness as independent variable and A’ as dependent variable also revealed a partial mediation effect (total effect: β = 0.27, *p* = 0.001; indirect effect: β = 0.11, *p* = 0.004; Table 4, Figure 3). This suggests that the SAME memory score mediated 40.74% of the total effect between Openness and A’.

**Figure 3.**
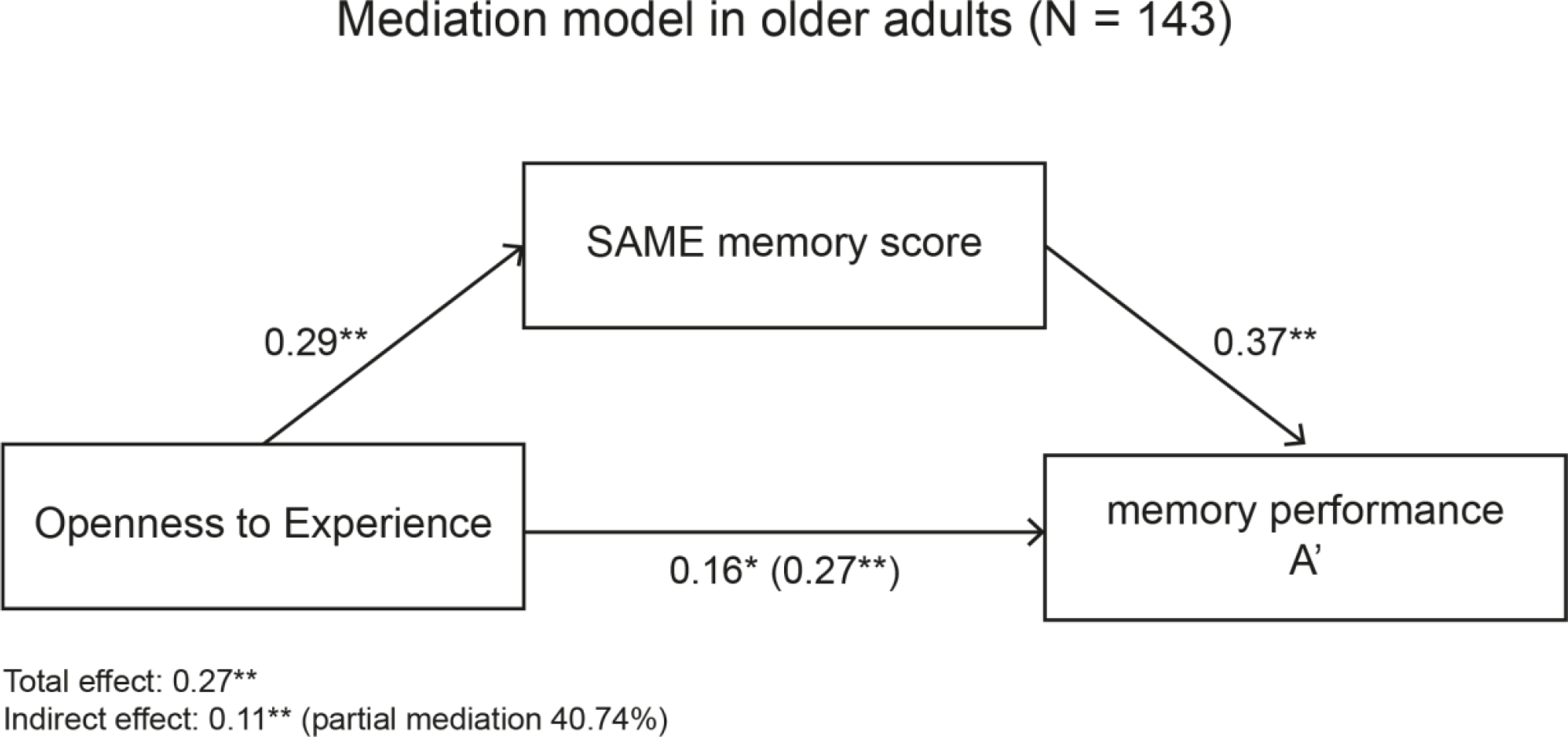
Mediation analysis within the group of older adults shows that the relationship between trait Openness and the memory performance (A’) is partly mediated (40.74% of the total effect) by the SAME memory fMRI score. \**p* < 0.05; \*\**p* < 0.01.

### Control analyses on the associations between Openness, memory performance, and fMRI measures

To ensure the observed mediation effects were not due to covariations with demographics and crystallized intelligence, the analyses described above were also conducted for Openness, A’, and FADE/SAME memory scores adjusted for age, gender, education, and MWT-B scores. We were able to confirm the partial mediation effects in these additional analyses (supplementary material B).

Since Extraversion showed a similar correlational pattern with memory performance and fMRI scores as Openness, mediation analyses including Extraversion instead of Openness were calculated, which did not reach statistical significance (supplementary material C). Moreover, partial correlations between Openness and A’ and FADE/SAME memory scores were still statistically significant after controlling for Extraversion (0.22 ≤ |*r*s| ≤ 0.25, *p*s < 0.001), while Extraversion (controlled for Openness) was only significantly correlated to A’ (*r* = 0.20, *p* = 0.019) but not with FADE/SAME memory scores (FADE: *r* = -0.11; SAME: *r* = 0.10; *p*s ≥ 0.204; supplementary material D1). A mediation analysis including Openness (controlled for Extraversion), A’ and fMRI memory scores revealed full mediation effects for FADE (indirect effect 33.33% of the total effect) and SAME memory scores (47.52%; supplementary material D2).

Additionally, trait theories suggest a meta-trait Plasticity, which is defined as the shared variance of Openness and Extraversion (DeYoung, 2006; DeYoung, 2013; DeYoung, 2015). We further run mediation analyses with the latent factor Plasticity as independent variable (supplementary material D3). These analyses revealed a significant indirect effect with the SAME memory scores (21.20% of total effect), but not the FADE memory scores (11.81% of total effect) as mediation variables.

### Voxel-wise association of fMRI contrasts with Openness

As an exploratory analysis, we computed voxel-wise regressions of the fMRI memory contrast with Openness in older adults. We observed significantly positive associations of Openness with successful memory encoding activations in left (β = 0.21, SD = 0.20*; t*(141) = 4.35, *p* = 0.009, × y z = -42 -79 23; 47 voxels) and right (β = 0.08, SD = 0.16*; t*(141) = 4.15, *p* = 0.025, × y z = 33 -79 2; 38 voxels) medial occipital gyrus (MOG; FWE cluster-level correction; Figure 4A). Additionally, the analysis revealed a positive association with the precuneus/posterior cingulate cortex (β = 0.14, SD = 0.25), which however did not reach corrected statistical significance (*t*(141) = 4.50, *p* = 0.080, × y z = 15 -58 17, 28 voxels; FWE cluster-level correction; supplementary material E, Figure S1).

**Figure 4.**
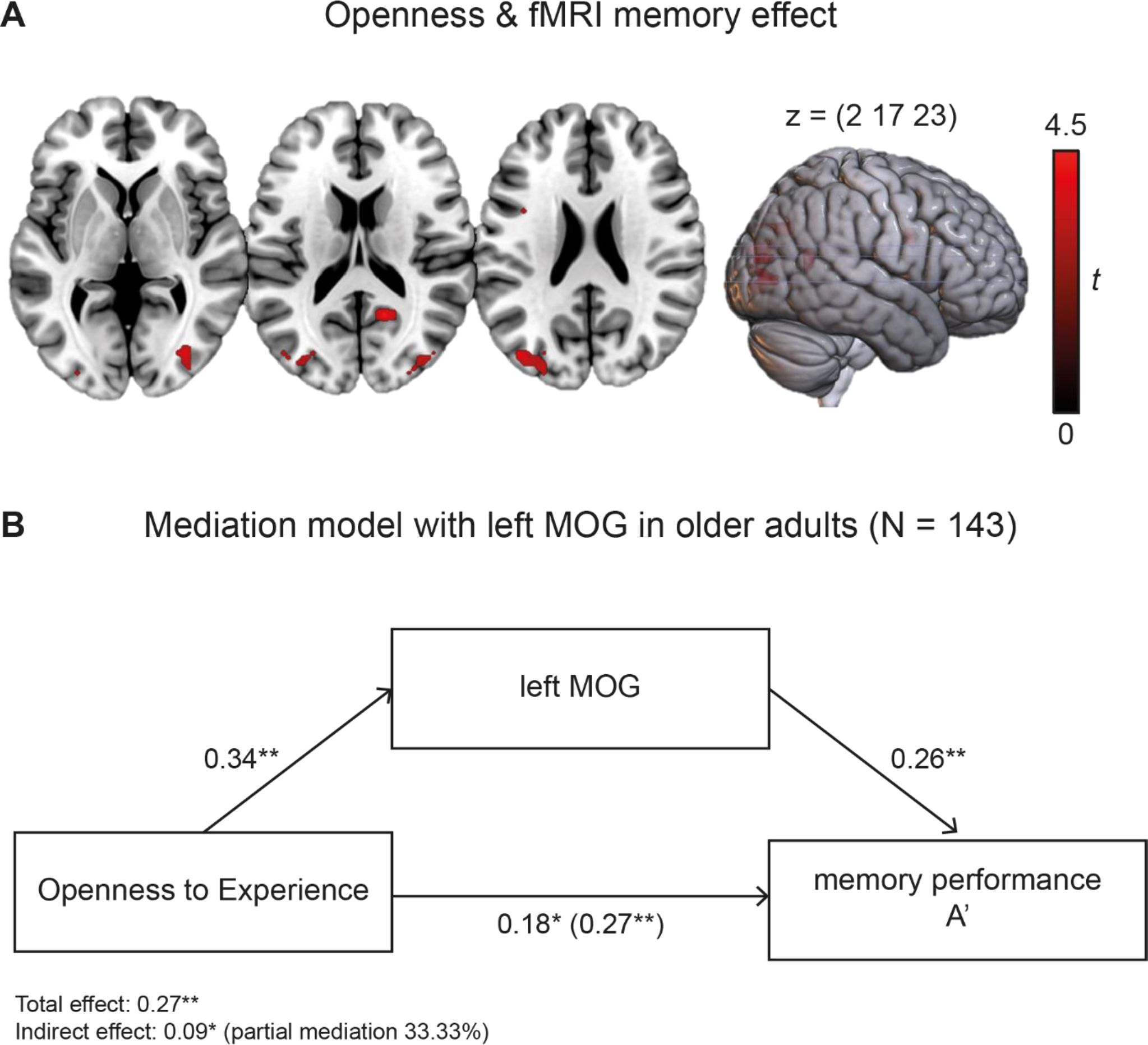
(A) Regression analysis of Openness and fMRI memory effect (positive effect) in older adults. *p* < 0.05, FWE-corrected at cluster level, cluster-defining threshold *p* < 0.001, uncorrected. All activation maps are superimposed on the MNI template brain provided by MRIcroGL (https://www.nitrc.org/projects/mricrogl/). (B) Mediation analysis showed that the relationship between Openness and memory performance (A’) was partially mediated (33.33% of the total effect) by activity in the left MOG. \**p* < 0.05; \*\**p* < 0.01.

Correlations between the left and right MOG peak voxel activity and personality traits as well as the FADE/SAME memory scores and A’ are shown in Table 5. A mediation analysis including Openness as independent, A’ as dependent, and left MOG peak voxel activity as mediation variable revealed a partial mediation (total effect: β = 0.27, *p* = 0.001; indirect effect: β = 0.09, *p* = 0.01, 33.33% of the total effect; Figure 4). Openness was positively related to A’ (β = 0.18, *p* = 0.03), and left MOG (β = 0.34, *p* < 0.001). There was a significant positive association between left MOG and A’ (β = 0.26, *p* = 0.002).<colcnt=1>

**Table 5.**
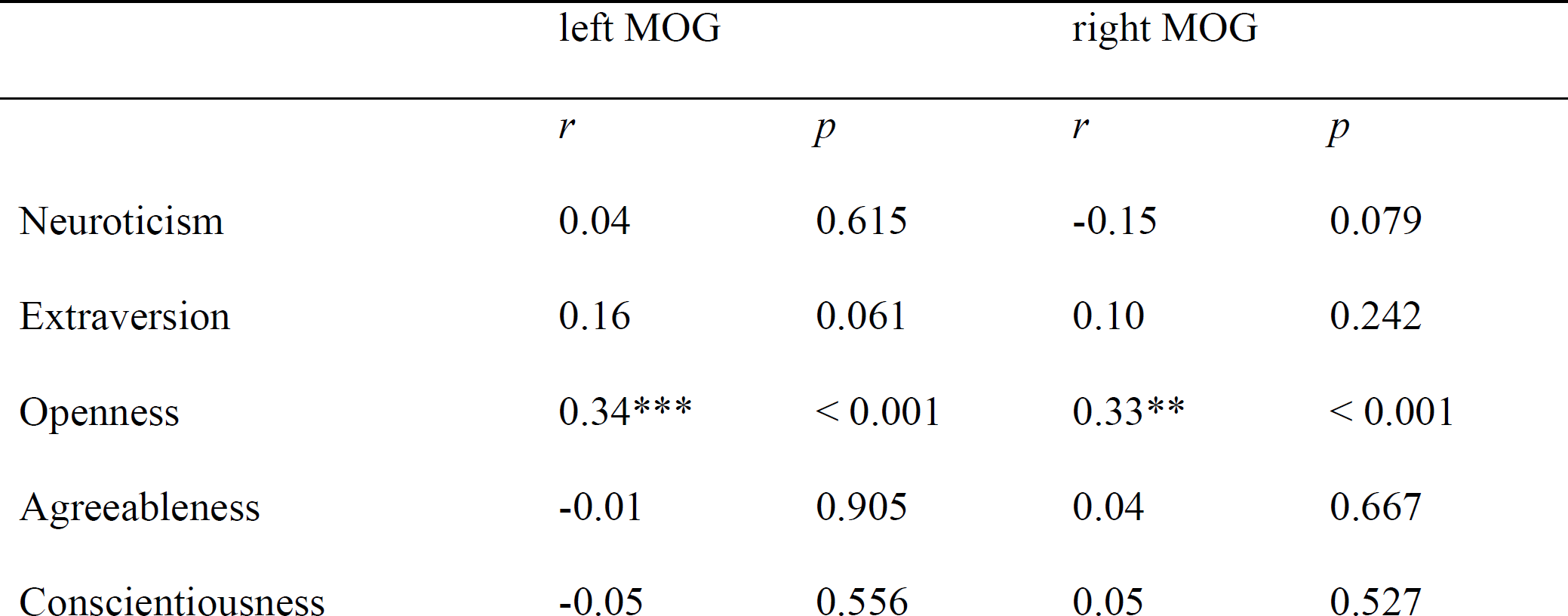
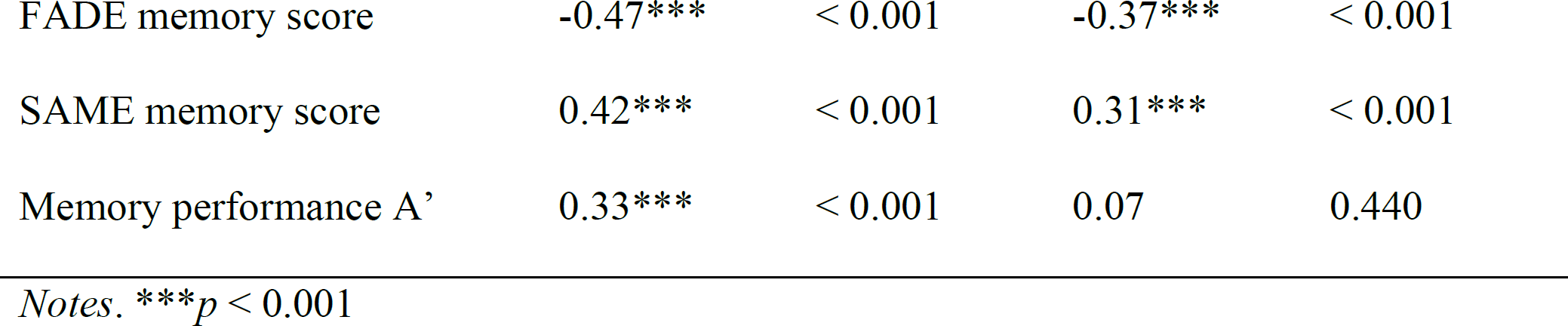
Pearson’s correlations between occipital gyrus peak-voxel activity (left & right MOG), Big Five personality traits, and memory-related brain activity and performance.

|  | left MOG |  | right MOG |  |
| --- | --- | --- | --- | --- |
|  | *r* | *p* | *r* | *p* |
| Neuroticism | 0.04 | 0.615 | -0.15 | 0.079 |
| Extraversion | 0.16 | 0.061 | 0.10 | 0.242 |
| Openness | 0.34*** | < 0.001 | 0.33** | < 0.001 |
| Agreeableness | -0.01 | 0.905 | 0.04 | 0.667 |
| Conscientiousness | -0.05 | 0.556 | 0.05 | 0.527 |
| FADE memory score | -0.47*** | < 0.001 | -0.37*** | < 0.001 |
| SAME memory score | 0.42*** | < 0.001 | 0.31*** | < 0.001 |
| Memory performance A' | 0.33*** | < 0.001 | 0.07 | 0.440 |

Notes. \*\*\* $p < 0.001$

## Discussion

The goal of the present study was to investigate to what extent differences in memory encoding-related brain activity explain the frequently reported relationship between Openness and memory performance in older adults. To this end, trait Openness was correlated with single-value fMRI scores reflecting deviations from or similarities with prototypical memory encoding brain activity in a visual memory encoding task. Consistent with our hypotheses, high trait Openness in older adults was related to both, better memory performance and more prototypical brain activity (low FADE/high SAME score). A mediation analysis revealed that the relationship between Openness and memory performance in older adults was substantially mediated by FADE/SAME fMRI scores. This overall provides first evidence that older adults scoring high on trait Openness show better memory performance, which is mediated by a more prototypical memory brain-network activity during encoding.

### Mediation of Openness and episodic memory performance by neural activity

Across all subjects, age group (older vs. young adults) had a significant effect on episodic memory performance and memory-related fMRI activity as indicated by FADE and SAME scores. This converges with previous studies demonstrating age differences in memory performance and functional brain activity (Düzel *et al*., 2011; Maillet and Rajah, 2014; Fjell *et al*., 2016; Damoiseaux, 2017; Morcom and Henson, 2018; Soch, Richter, Schütze, Kizilirmak, Assmann, Behnisch, *et al*., 2021; Soch, Richter, Schütze, Kizilirmak, Assmann, Knopf, *et al*., 2021).

The link between Openness and episodic memory retrieval in older adults was mediated by activity in memory brain networks during memory encoding. Specifically, older adults who scored high in trait Openness also showed more prototypical brain activity during memory encoding, which was directly related to better episodic memory performance. These results converge with studies reporting a positive relationship between Openness and episodic memory performance in older adults (Terry *et al*., 2013; Luchetti *et al*., 2016; Sutin *et al*., 2019; Stephan *et al*., 2020). One possible explanation for this may be that highly open older adults are more likely to engage into intellectually stimulating activities such as travelling, attending cultural events, seeking novel or aesthetic experiences (Kraaykamp and van Eijck, 2005; Chamorro-Premuzic and Furnham, 2008; Soubelet and Salthouse, 2010; Jackson *et al*., 2020; Talpain and Soubelet, 2022). Increased exposure to cognitive-stimulating life experiences may delay age-related manifestation of memory deficits (Stern, 2009; Franchow *et al*., 2013). Additionally, Openness is related to a broad range of cognitive abilities, including fluid and crystallized intelligence, and working memory capacity (Gregory *et al*., 2010; Schretlen *et al*., 2010; Sharp *et al*., 2010; Chapman *et al*., 2012; Nishita *et al*., 2019). In this context, high Openness may reflect a protective factor at personality trait level, indicating preserved memory-related cognitive functioning in older age.

At structural brain levels, there are overlapping reports that high Openness is related to a decrease in cortical thickness particularly in frontal (orbitofrontal, superior and posterior frontal, and prefrontal) and parietal (inferior parietal and precuneus) regions (Wright *et al*., 2007; Kapogiannis *et al*., 2013; Vartanian *et al*., 2018). Other structural measures linked high Openness to increased orbitofrontal cortex folding and surface area (Vartanian *et al*., 2018) and a lower integration of structural brain networks (Talaei and Ghaderi, 2022). In contrast, other studies found small-to-no associations between Openness and structural brain measures (DeYoung *et al*., 2010; Bjørnebekk *et al*., 2013; Avinun *et al*., 2020; Hyatt *et al*., 2022). These null findings were further supported by recent meta-analyses that did not discover any replicable relationship between Openness (and other Big Five traits) and cortical thickness, gray matter volume, and surface areas (Chen and Canli, 2022). At functional levels, the neural basis of Openness has largely been associated with resting state fMRI activity. These studies showed that individual differences in Openness can be predicted by functional connectivity matrices (Dubois *et al*., 2018) and have repeatedly been related to activity in the DMN (Blain et al., 2020; Horn et al., 2014), suggesting highly Open individuals show a more efficient DMN functioning (Beaty *et al*., 2016; Beaty *et al*., 2018). Another recent study reported a decreased within-network DMN functional connectivity in highly Open individuals, which was mainly driven by the Fantasy sub-facet of Openness (Marstrand-Joergensen *et al*., 2021). DMN activity also was related to cognitive functions with particular relevance for trait Openness (e.g., creativity, imagination, & divergent thinking; DeYoung, 2014; DeYoung et al., 2012) and episodic memory retrieval. The DMN is subject to age-related changes, indicating alterations in the DMN such as reduced activity in medial-temporal and parietal regions and increased activity in frontal areas, which often have been assumed to reflect compensatory reactions to age-related brain alterations (Desgranges *et al*., 2011; Andrews-Hanna *et al*., 2014; Mohan *et al*., 2016; Mak *et al*., 2017). The mentioned regions in the DMN also have a qualitative overlap with brain regions whose activity was measured using FADE/SAME memory scores, such as the precuneus and posterior cingulate cortex. Overall, it may be that high Openness reflects a protective factor when it comes to successful aging of neural structures associated with episodic memory functioning. In the present study, this might be reflected by a higher similarity of highly open older adults’ brain activity to the prototypical functional brain responses during memory encoding.

#### Exploratory analysis: Associations between Openness and medial occipital brain activity

In an exploratory, voxel-wise analysis within older adults, we found that high Openness and better memory performance were related to amplified activations in the MOG. Converging with our results, the MOG shows robust activations during encoding (Maillet and Rajah, 2014) and recognition (Yonelinas *et al*., 2001) as well as increased functional connectivity with the hippocampus during successful memory formation (Ranganath *et al*., 2005). Hence, the MOG is considered part of a visual memory and perception network, which is active during both episodic memory encoding and retrieval (Slotnick, 2004; Slotnick and Schacter, 2006; Thakral and Slotnick, 2015; Waldhauser *et al*., 2016). This is in line with a proposed model for visual memory encoding, which demonstrated an early involvement of the MOG driven by the primary and secondary visual areas (Nenert *et al*., 2014). These areas are also affected by aging, indicating decreased occipital activity during working memory, visual attention, and episodic retrieval tasks (Cabeza *et al*., 2004; Davis *et al*., 2008), and a disproportionate recruitment of left vs. right visual cortex during encoding of visual episodic information (Maillet and Rajah, 2014). There is preliminary evidence suggesting that high Openness is related to different low-level visual perceptual experiences (Antinori *et al*., 2017). Accordingly, Blake & Palmisano (2021) reported differences in perceptual processing of ambiguous visual stimuli in high creativity/divergent thinking, two features which are also related to individual differences in Openness (DeYoung *et al*., 2012; Kaufman *et al*., 2016). This overall raises the question whether the age-related associations between Openness and successful memory encoding may not only rely on preserving neurocognitive functioning by intellectual engagement and activities, but also on personality-related differences in early visual processing during memory encoding.

### Clinical relevance

In a recent meta-analysis, AD patients showed lower levels of self-reported and informant-obtained Openness and Extraversion, and higher levels of Neuroticism compared to controls (D’Iorio *et al*., 2018). A similar personality profile was considered a risk factor and predictor for developing AD (Caselli et al. 2018; Duberstein *et al*., 2011) and assumed to change before vs. after the AD diagnosis (Robins Wahlin and Byrne, 2011). Other studies proposed that specifically low Openness was a pre-clinical marker of incipient cognitive decline (Williams *et al*., 2013), and that high Openness may protect from memory decline (Terry *et al*., 2013). This suggests that Openness is part of a personality profile in which high Openness may constitute a protective factor in the transition from healthy memory functioning to risk-stages for AD and personality changes that go along the development of AD. Our study is thus of high clinical relevance, as it provides novel evidence that the presumably protective effect of Openness on episodic memory functioning is reflected by functional brain-network differences. This should encourage conducting similar studies in individuals at risk of developing AD to better understand the beneficial characteristics of personality traits on brain alterations before and after AD onset.

#### A potential role for the mesolimbic dopamine system and Openness in preserved cognition

In the present study, memory network activity and performance in older adults were both specifically correlated with Openness and Extraversion, which may potentially emphasize the influence of the dopamine system. This converges with empirical and theoretical references that ascribe dopaminergic functioning a major role in the cognitive and behavioral expressions of Openness and Extraversion (Depue and Lenzenweger, 2001; DeYoung, 2010; DeYoung, 2013; Pickering and Pesola, 2014; Wacker and Smillie, 2015). Extraversion has previously been linked to dopamine-rich brain regions (Li *et al*., 2019) and cortical reward processing (Cooper *et al*., 2014; Smillie *et al*., 2019). In an fMRI study, Passamonti et al. (2015) reliably linked high Openness to increased functional connectivity of prefrontal cortex and substantia nigra/ventral tegmental area. Moreover, a recent pharmacological study linked Openness and divergent thinking to dopamine activity (Käckenmester *et al*., 2019). Furthermore, in the cybernetic big five theory, Openness and Extraversion are considered a meta-trait Plasticity whose neuromodulator is dopamine (DeYoung *et al*., 2005; DeYoung, 2013; DeYoung, 2015). Our analyses including Plasticity (supplementary material D) indicated a mediation effect solely for the SAME memory score, which may suggest an involvement of dopamine. Dopamine has also often been implicated in episodic memory functioning (Wittmann *et al*., 2005; Adcock *et al*., 2006; Düzel *et al*., 2010), and impaired dopaminergic neurotransmission is associated with cognitive aging and longitudinal memory decline (Bäckman and Farde, 2009; Morcom *et al*., 2010; Chowdhury *et al*., 2012; Papenberg *et al*., 2014; Nyberg *et al*., 2016). Düzel et al. (2010) proposed that an increase of mesolimbic dopaminergic activity promotes exploratory behavior and ultimately memory performance in older adults. In line with this framework, our results suggest links between Openness/Extraversion and age-related differences in neural memory functioning.

### Relevance of Openness facets

A major limitation of the present study was that personality traits were assessed with only one questionnaire (NEO-FFI), which ultimately did not allow to examine by which trait sub-facet the age-related Openness effects on memory were mainly driven. Accordingly, previous studies reported that Openness facets (e.g., fantasy, feelings, ideas) had different longitudinal trajectories (Bleidorn *et al*., 2009) and varying associations with psychometric intelligence (Moutafi *et al*., 2006). Other studies that divided Openness into the sub-facets “Openness” (fantasy, aesthetics) and “Intellect” (intellectual engagement; DeYoung, 2014) linked Intellect to creative achievements in the sciences (Kaufman *et al*., 2016), fluid intelligence (Nusbaum and Silvia, 2011), and increased working memory-related prefrontal activity (DeYoung *et al*., 2009). In contrast, the sub-facet Openness, but not Intellect was related to implicit learning (Kaufman *et al*., 2010) and reduced cortical thickness in fronto-temporal gyri (Vartanian *et al*., 2018). In the development of the Big Five Aspects Scale (BFAS; DeYoung, 2013; DeYoung *et al*., 2013; DeYoung, 2014), the sub-facets Openness and Intellect were correlated with the NEO-PI-R Openness scale, indicating that NEO Openness total scores may stronger reflect the BFAS sub-facet Openness than Intellect (DeYoung *et al*., 2007). An exploratory analysis of single-item correlations (see supplementary material A) showed that memory performance was particularly related to an item of the NEO-PI-R Ideas facet (i.e., open-mindedness, open to new ideas, intellectual engagement), and FADE/SAME memory scores to an item of the Aesthetics facet (i.e., open to and interests in art, music, and poetry). According to the independent associations of Openness sub-facets with individual differences at cognitive, behavioral, and neural levels, it seems very encouraging for follow-up studies to assess personality traits with a higher resolution (e.g., NEO-PI-R or BFAS; DeYoung, 2007). Such studies would not only provide a better understanding of the Openness sub-facets, but would also shed more light on the specific cognitive processes (e.g., creativity vs. intellect) involved in the protective effects of Openness on neurocognitive aging. In the context of the present study, it might be conceivable that the effects of Openness on cortical memory encoding and memory performance may be even stronger represented in one of the facets than the global Openness domain.

### Specificity of trait activation

The associations of Openness with neural memory encoding were only supported in the older adult age group. This might be due to group-related differences in average Openness and memory performance. In line with other research, in the present study, older vs. young adults showed lower Openness scores (Schaie *et al*., 2004; Roberts *et al*., 2006; Allemand *et al*., 2007; Bleidorn *et al*., 2009) and a declined memory performance (Oschwald *et al*., 2019). It is conceivable that the memory task was not challenging enough for young adults, such that it did not pose an optimal situation to activate the influences of personality traits (McNaughton and Smillie, 2018; Taconnat *et al*., 2022) and thus masked potential shared variance between memory encoding and trait Openness. Moreover, the FADE/SAME scores are by definition contrasted with the memory-related fMRI activity of the young group. Thereby, the FADE/SAME scores might further conceal potential trait correlations within the group of young adults and rather reflect highly suitable single-scores for the investigation of personality in neurocognitive aging.

### Conclusion

Our study provides novel evidence that the personality trait Openness is associated with episodic memory performance and this effect was substantially mediated by memory-related brain activity in older adults. Specifically, those older adults with high Openness showed better memory performance and this was due to a higher similarity to brain activity patterns of young adults. To the best knowledge of the authors, this is the first report linking the relationship between trait Openness and memory performance to actual brain activity in an episodic memory task. The present study further shows the utility of single-value fMRI scores in personality neuroscience with a high statistical power.

## Statements

## Acknowledgments

The authors would like to thank Hartmut Schütze for programming the fMRI paradigm. We thank Anne Assmann, Adriana Barman, Hannah Feldhoff, Larissa Fischer, Marieke Klein, Lea Knopf, Matthias Raschick, Annika Schult for help with behavioral data collection. We are grateful to Kerstin Möhring, Ilona Wiedenhöft, Katja Neumann, and Claus Tempelmann for assistance with MRI data acquisition.

## Data Availability Statement

Due to data protection regulations, sharing of the entire data set underlying this study in a public repository is not possible. We have previously provided GLM contrast images as a NeuroVault collection (https://neurovault.org/collections/QBHNSRVW/) and MATLAB code for imaging scores as a GitHub repository (https://github.com/JoramSoch/FADE_SAME) for an earlier article using the same dataset (Soch, Richter, Schütze, Kizilirmak, Assmann, Behnisch, et al., 2021). Access to de-identified raw data will be provided by the authors upon reasonable request.

## Footnotes

1 The MWT-B was assessed for 142 of the n= 143 older adults and 93 of the n = 209 young adults.

