## Supplementary Material for "Openness to experience is associated with neural and performance measures of memory in older adults"

Running title: Openness in memory-related neural aging

Christopher Stolz[1,2*], Ariane Bulla[1], Joram Soch[3,4], Björn H. Schott[1,3,5], Anni Richter[1,6*]

[1]Leibniz Institute for Neurobiology (LIN), Magdeburg, Germany

[2]Institute of Psychology, Otto-von-Guericke University Magdeburg, Magdeburg, Germany

[3]German Center for Neurodegenerative Diseases (DZNE), Göttingen, Germany

[4]Bernstein Center for Computational Neuroscience (BCCN), Berlin, Germany

[5]Department of Psychiatry and Psychotherapy, University Medical Center Göttingen, Göttingen, Germany

[6]Center for Intervention and Research on adaptive and maladaptive brain Circuits underlying mental health (C-I-R-C), Jena-Magdeburg-Halle

***Correspondence**

Anni Richter, Leibniz Institute for Neurobiology, Department of Behavioral Neurology, Brenneckestraße 6, D-39118 Magdeburg, Germany.. Telephone: 0049-391-626393151.

Christopher Stolz, Otto-von-Guericke University, Department of Neuropsychology Universitätsplatz 2, D-39106 Magdeburg, Germany.

### Supplementary Material A: Single-item correlations of NEO-FFI Openness with memory performance and FADE/SAME memory scores

Table S1 shows the correlations between the NEO-FFI Openness items and the memory performance (A’) and FADE/SAME memory scores. The largest correlation of an Openness item with the respective memory measures are denoted in bold. The largest correlation for A’ was found for O2 “Philosophical discussions (R)” which belongs to the NEO-PI-R Openness facet Ideas (open-mindedness, open to new ideas, intellectual engagement). For FADE/SAME memory scores the largest correlations were found for item O9 “Wave of excitement to art/literature” which belongs to the NEO-PI-R Openness facet Aesthetics (open to and interests in art, music, and poetry). In comparison with the Openness/Intellect taxonomy of trait Openness measured with the Big Five Aspects Scale (BFAS; DeYoung, 2013; DeYoung *et al.*, 2013; DeYoung, 2014), the NEO-PI-R facets of Openness were particularly related to the BFAS Openness facet (DeYoung *et al.*, 2007), suggesting that Openness total scores assessed via NEO-PI-R/NEO-FFI may stronger reflect BFAS Openness than BFAS Intellect.

**Table S1**

Correlations between single items of NEO-FFI trait Openness and memory performance and FADE/SAME memory scores.

| NEO-FFI Openness item | NEO-PI-R Facet | A’ | FADE memory score | SAME memory score |
|---|---|---|---|---|
| O1 Daydreaming (R) | Fantasy | -0.02 | -0.01 | -0.01 |
| **O2 Philosophical discussions (R)** | **Ideas** | **0.25**** | -0.12 | 0.16 |
| O3 Patterns in arts/nature | Aesthetics | 0.13 | -0.03 | 0.14 |
| 04 Controversial speakers (R) | Values | 0.12 | -0.14 | 0.21* |
| 05 Impression of poetry (R) | Aesthetics | 0.17* | -0.12 | 0.20* |

| | | | | |
|---|---|---|---|---|
| 06 New and foreign foods | Actions | 0.12 | -0.11 | 0.14 |
| O7 Notice moods (R) | Feelings | 0.18* | -0.15 | 0.17 |
| O8 Respect religious authorities | Values | <0.01 | -0.03 | -0.08 |
| **O9 Wave of excitement to art/literature** | **Aesthetics** | 0.14 | **-0.25**** | **0.23**** |
| 010 Interest in speculating (R) | Ideas | 0.14 | -0.06 | 0.13 |
| O11 Curiosity | Ideas | 0.05 | -0.13 | 0.10 |
| O12 Enjoy theories | Ideas | 0.15 | -0.21* | 0.20* |

Notes. R = reversed

### Supplementary Material B: Mediation analysis with residual variables adjusted by age, gender, education, and MWT-B

Multiple regression analyses were conducted including age, gender, education, and MWT-B scores on Openness, A', FADE memory score, and SAME memory score and residuals were saved (i.e. adjusted for demographics and the MWT-B) for further mediation analyses.

The mediation analysis with the FADE memory score as mediator on the relationship between Openness as independent variable and A' as dependent variable revealed a partial mediation effect (total effect: $\beta = 0.24$, $p = 0.003$; indirect effect: $\beta = 0.06$, $p = 0.041$). Openness still had a significant positive relationship with A' ($\beta = 0.18$, $p = 0.023$), and was negatively related to the FADE memory score ($\beta = -0.22$, $p = 0.007$, that is higher Openness was associated with a lower brain network deviation. There was a significant negative association between the FADE memory score and A' ($\beta = -0.25$, $p = 0.002$). Overall, this suggests that 25.10% of the total effect between Openness and A' was mediated by the FADE memory score.

The mediation analysis with the SAME memory score as mediator on the relationship between Openness as independent variable and A' as dependent variable revealed a full mediation (total effect: $\beta = 0.24$, $p = 0.003$; indirect effect: $\beta = 0.09$, $p = 0.010$). In the mediation model, Openness was not significantly related to A' anymore ($\beta = 0.15$, $p = 0.057$). Openness was significantly related to the SAME memory score ($\beta = 0.26$, $p = 0.001$), that is higher Openness was associated with an increased brain network similarity. There was a significant positive association between SAME memory score and A', $\beta = 0.34$, $p < 0.001$. Overall, this suggests that 37.65% of the total effect between Openness and A' was mediated by the SAME memory score.

### Supplementary Material C: Mediation analysis with Extraversion, SAME/FADE memory scores, and memory performance

The results of mediation analyses with FADE/SAME memory scores as mediator on the relationship between Extraversion and memory performance (A') are presented in Table S2. Overall, these results suggest relatively small indirect effects, which did not reach statistical significance, and thus do not provide empirical evidence for a partial mediation effect of FADE/SAME memory scores on the relationship between Extraversion and memory performance.

**Table S2**

Mediation analysis within the group of older adults with FADE/SAME memory scores as mediators of the association between Extraversion and episodic memory performance (A').

| **FADE memory score** | β | *p* |
|---|---|---|
| Total effect: Extraversion → A' | 0.25 | 0.001 |
| Direct effect: FADE memory score → A' | -0.30 | < 0.001 |
| Direct effect: Extraversion → A' | 0.20 | 0.009 |
| Indirect effect: Extraversion → FADE memory score → A' | 0.05 (19.44%) | 0.077 |
| **SAME memory score** | β | *p* |
| Total effect: Extraversion → A' | 0.25 | 0.004 |
| Direct effect: SAME memory score → A' | 0.38 | < 0.001 |
| Direct effect: Extraversion → A' | 0.19 | 0.013 |
| Indirect effect: Extraversion → SAME memory score → A' | 0.06 (25.40%) | 0.059 |

### Supplementary Material D: Exploratory analysis including trait Openness and Extraversion

#### D1: Partial correlations between Openness/Extraversion and memory measures

The Table S3 shows the partial correlations between Openness (controlled for Extraversion) and Extraversion (controlled for Openness) and the memory performance (A') and FADE/SAME memory scores. Openness controlled for Extraversion is still significantly correlated to the performance and fMRI scores. Extraversion controlled for Openness is positively associated with the memory performance but there is no significant relationship with the FADE/SAME memory scores.

**Table S3**

Partial correlations between Openness (controlled for Extraversion) and Extraversion (controlled for Openness) and memory performance (A') and FADE/SAME memory scores.

| | Openness w/o Extraversion | Extraversion w/o Openness |
|---|---|---|
| Memory performance (A') | 0.22** | 0.20* |
| FADE memory score | -0.22** | -0.11 |
| SAME memory score | 0.25** | 0.10 |

#### D2: Mediation analysis with Openness controlled for Extraversion

The results of mediation analyses with FADE/SAME memory scores as mediator on the relationship between Openness controlled for Extraversion and memory performance (A') are presented in Table S4. Overall, these analyses replicate the result pattern of the study's central findings, but extends them with the findings of a full mediation for both mediator variables, FADE and SAME memory scores.

**Table S4**

Mediation analysis within the group of older adults with FADE/SAME memory scores as mediators of the association between Openness w/o Extraversion and episodic memory performance (A').

| **FADE memory score** | β | *p* |
|---|---|---|
| Total effect: Openness w/o Extra. → A' | 0.21 | 0.009 |
| Direct effect: FADE memory score → A' | -0.30 | < 0.001 |
| Direct effect: Openness w/o Extra. → A' | 0.15 | 0.065 |
| Indirect effect: Openness w/o Extra. → FADE memory score → A' | 0.07 (33.33%) | 0.030 |
| **SAME memory score** | β | *p* |
| Total effect: Openness w/o Extra. → A' | 0.21 | 0.009 |
| Direct effect: SAME memory score → A' | 0.38 | < 0.001 |
| Direct effect: Openness w/o Extra. → A' | 0.12 | 0.143 |
| Indirect effect: Openness w/o Extra. → SAME memory score → A' | 0.10 (47.52%) | 0.008 |

**D3: Mediation analysis with meta-trait Plasticity (shared variance of Openness and Extraversion), SAME/FADE memory scores, and memory performance**

The results of mediation analyses with FADE/SAME memory scores as mediator on the relationship between Plasticity (latent factor, shared variance of Openness and Extraversion) and memory performance (A') are presented in Table S5.

**Table S5**

Mediation analysis within the group of older adults with FADE/SAME memory scores as mediators of the association between Plasticity and episodic memory performance (A’).

| **FADE memory score** | β | *p* |
|---|---|---|
| Openness → Plasticity | 0.56 | - |
| Extraversion → Plasticity | 0.46 | 0.005 |
| Total effect: Plasticity → A’ | 0.51 | 0.011 |
| Direct effect: FADE memory score → A’ | -0.14 | 0.218 |
| Direct effect: Plasticity → A’ | 0.45 | 0.035 |
| Indirect effect: Plasticity → FADE memory score → A’ | 0.06 (11.81%) | 0.146 |
| **SAME memory score** | β | *p* |
| Openness → Plasticity | 0.58 | - |
| Extraversion → Plasticity | 0.44 | 0.005 |
| Total effect: Plasticity → A’ | 0.50 | 0.012 |
| Direct effect: SAME memory score → A’ | 0.23 | 0.052 |
| Direct effect: Plasticity → A’ | 0.39 | 0.054 |
| Indirect effect: Plasticity → SAME memory score → A’ | 0.11 (21.20%) | 0.029 |

Overall, these results suggest a relatively small but significant mediation effect of SAME memory scores on the relationship between Plasticity and memory performance. The mediation analysis including FADE memory scores as mediator variable did not reach statistical significance.

### Supplementary Material E: Voxel-wise association of fMRI contrasts with Openness (Precuneus)

Additionally, voxel-wise analyses revealed a hint for an association with the precuneus/posterior cingulate cortex (β = 0.14, SD = 0.25) which however did not reach statistical significance ($t(141) = 4.50$, $p = 0.080$, x y z = 15 -58 17, 28 voxels; FWE cluster-level correction; Figure S1).

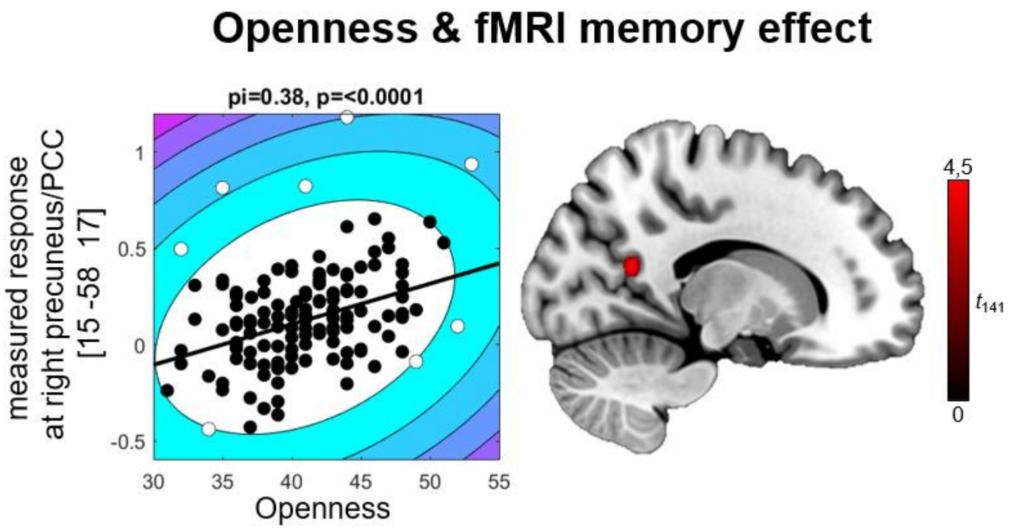


**Figure S1**. Regression analysis of Openness and fMRI memory effect (positive effect) in older adults. $p < 0.05$, FWE-corrected at cluster level, cluster-defining threshold $p < 0.001$, uncorrected. All activation maps are superimposed on the MNI template brain provided by MRIcroGL (https://www.nitrc.org/projects/mricrogl/). Correlational plots are outlier robust Shepherds Pi correlations (Samuel Schwarzkopf *et al.*, 2012) of peak voxel estimates with Openness.
